# Peptides derived from gp43, the most antigenic protein from *Paracoccidioides brasiliensis*, form amyloid fibrils *in vitro*: implications for vaccine development

**DOI:** 10.1101/2021.07.08.451661

**Authors:** Thyago R. Cardim-Pires, Ricardo Sant’Anna, Debora Foguel

## Abstract

Fungal infection is an important public health problem afflicting more than a billion people worldwide. Mycoses are especially important in Latin America, and in Brazil in particular. *Paracoccidioides* is the genus of fungi responsible for paracoccidioidomycosis comprising two species, *P. brasiliensis* and *P. lutzii*. The lungs are the primary infection site, but oral mucosa and airways can also be affected. The glycoprotein gp43 is involved in fungi adhesion to epithelial cells and is the most studied protein of *P. brasiliensis*. Seminal work identified a specific stretch of 15 amino acids that spans the region 181-195 (called P10) as an important epitope of gp43, being recognized by T lymphocytes in peripheral blood mononuclear cells of mice and humans and is envisioned as a potential vaccine component. Here, we show by using thioflavin T (ThT), transmission electron microscopy and other methods that synthetic P10 forms typical amyloid aggregates in solution in very short times, a property that could hamper vaccine development. *In silico*, aggregation analysis reveals several aggregation-prone regions (APR) in the P10 sequence that are capable of forming amyloid cores with steric zipper architecture. Seeds of P10 obtained by fibril mechanical fragmentation were able to induce the aggregation of P4 but not P23, as evidenced by ThT binding and mass spectrometry. These two peptides, also derived from gp43, are potent modulators of local and systemic inflammation. *In-silico* proteolysis studies with gp43 revealed that aggregation-prone, P10-like peptides could be generated by the action of several proteases such as proteinase K, trypsin and pepsin, which suggests that P10 could be formed upon gp43 digestion in a physiological condition. Considering our data in the context of a potential vaccine development, we redesigned the sequence of the P10 peptide, maintaining the antigenic region (HTLAIR), but drastically reducing its aggregation propensity.

## Introduction

Fungal infection is an important public -health problem afflicting more than a billion people and contributing to the death of 1-2% of the patients. Mycoses are especially important in Latin America, and in Brazil in particular (80% of the cases), where the climate favors yeast proliferation and infection, prevailing in rural workers of endemic areas. *Paracoccidioides* is the genus of fungi responsible for paracoccidioidomycosis (PCM) comprising two species, *P. brasiliensis* (Pb) and *P. lutzii*. The lungs are the primary infection site, but oral mucosa and airways can also be affected (GONZALEZ and HERNANDEZ, 2016).

The success of yeast-host interaction depends on several regulatory mechanisms as well as the expression of several different virulence factors. Among these are adhesins, which are surface proteins that recognize extracellular matrix (ECM) components of the host cells (SARDI et al., 2015).

Gp43 is a glycoprotein involved in fungi adhesion to epithelial cells and macrophages and is the most studied protein of *Paracoccidioides brasiliensis*, although other adhesins (moonlighting proteins) are also important to fungal virulence (SARDI et al., 2015; VICENTINI et al., 1994). Gp43 contains regions that drive interactions with laminin, collagen and fibronectin, allowing Pb adhesion to the cell (MENDES-GIANNINI et al., 2006). Gp43 has 416 amino acids, with secondary and tertiary structures not yet described. It presents high sequence homology (80% similarity) to the yeast exo-1,3-β-glucanase (UniProtKB -C1H4T0 **Figure 1A**), but no detected enzymatic activity (CISALPINO et al., 1996), since the catalytic site differs significantly in the Pb protein. Secreted during fungal infection, gp43 is the main antigen detected in patients (MARQUES DA SILVA et al., 2003) and it contains epitopes capable of eliciting a cellular immune response in animal models and in human patients leading to the production of IFN-γ by lymphocytes, which stimulates the formation of granulomas to contain the yeast cells (MAGALHÃES et al., 2011; KONNO et al., 2012).

**Figure 1.**
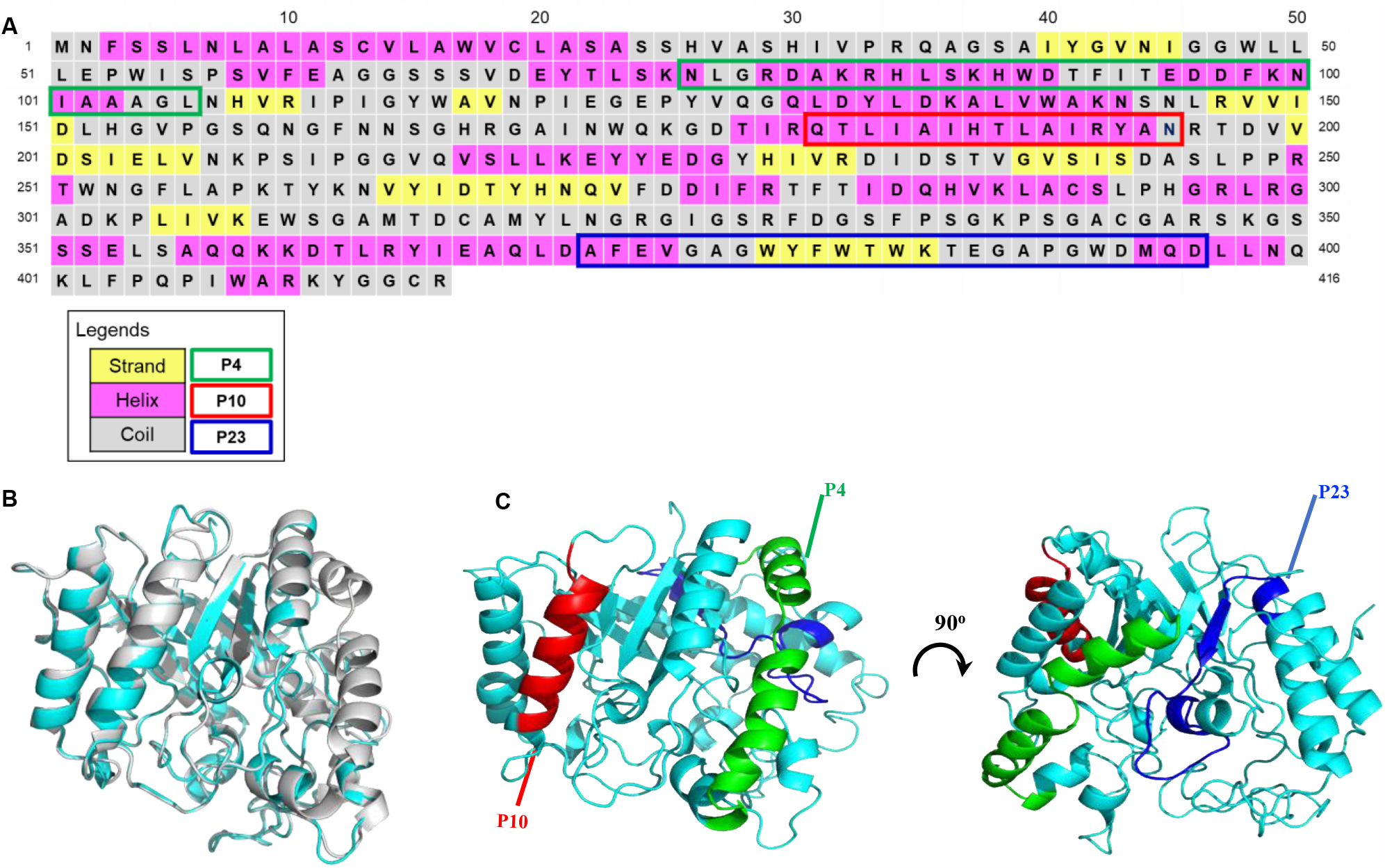
Primary sequence and predicted secondary structure elements of gp43 from *Paracoccidioides brasiliensis* strain B339 highlighting the three peptides studied here (P4, P10 and P23). (**A**) The primary structure was obtained at GenBank access number AAG36697.1. Secondary structure elements of gp43 protein were predicted by PSIPRED and are indicated by color Pink, yellow and gray cells indicate, respectively, α-helix, β-strand and random-coil structures. Green, red and blue outlines represent P4, P10 and P23 primary sequences, respectively. **(B)** The structural model of gp43 was generated by using RaptorX software (KÄLLBERG et al., 2012) using the β-glucanase structure as template. Gp43 is displayed in cyan and β-glucanase in gray. **(C)** Positions of P4, P10 and P23 peptides in the structural model of gp43 are depicted in green, red and blue, respectively. Superposition of both structures in B and C was generated by PyMOL (The PyMOL Molecular Graphics System, Version 1.2r3pre, Schrödinger, LLC.)

Since gp43 is the main PCM diagnostic antigen containing epitopes that elicit delayed-type hypersensitivity in animals and humans, in 1998, Taborda and coworkers synthesized 25 gp43-derived peptides in order to identify the T-cell epitopes. From this extensive study, the authors identified P10 (QTLIAIHTLAIRYAN), a specific stretch of 15 amino acids that spans the region 181-195 and was recognized by T lymphocytes in peripheral blood mononuclear cells of mice and humans (TABORDA et al., 1998; IWAI et al., 2003). Mice immunized with P10 developed lung infection 200 times less intense than the unimmunized mice, a response that was as effective as immunization with full-length gp43.

Later, Iwai and coworkers (2003) used an *in-silico* approach (TEPITOPE algorithm) (ZHANG et al., 2012) to localize peptide sequences from gp43 that would most likely bind multiple human leucocyte antigen-DR (HLA-DR) molecules and tested their recognition by T-cells from sensitized individuals. The most recognized peptide from peripheral blood mononuclear cells from PCM patients was gp43(180-194), recognized by T-cells from 53% of patients. This region of gp43 corresponds precisely to P10. Several groups are studying this peptide as a candidate for a vaccine against PCM and evaluating different adjuvants to potentiate the immunological response (MAYORGA et al., 2012). These studies include the use of a DNA vaccine encoding the P10 sequence (RITTNER et al., 2012). Magalhães and collaborators showed that dendritic cells primed with P10 protect the host against the development of mycosis and these cells are also effective in the treatment of well-established infections (MAGALHÃES et al., 2011).

The use of synthetic peptides to develop vaccines against microbes has been envisioned for a long time (SESARDIC, 1993). However, little is known about the biophysical properties of these peptides in solution, such as their tendency to aggregate into insoluble material, including amyloid fibrils. This latter type of aggregate is involved in amyloid diseases and small pieces of amyloid fibril can seed the aggregation of other cellular proteins, something unwanted for vaccine development (LI et al., 2014).

Here, we present evidence that synthetic P10 forms typical amyloid aggregates in solution in very short times (a few seconds). *In-silico* analyses using different algorithms revealed that gp43-derived peptides, including P10, have high aggregation propensity. Molecular modeling approaches showed that several stretches of the P10 sequence fit well into structural models of steric zippers, ensembles present in highly organized amyloid aggregates (EISENBERG et al., 2012). Interestingly, seeds formed by mechanical fragmentation of amyloid fibrils composed of P10 were able to induce the aggregation of a non-aggregating peptide also derived from gp43, named P4 (NLGRDAKRHLSKHWDTFITEDDFKNIAAAGL), but not the peptide P23 (AFEVGAGWYFWTWKTEGAPGWDMQD). Mass spectrometry showed that the fibrils composed of P4 seeded by P10 contains both sequences, suggesting a co-aggregational mechanism. P4 and P23 are potent modulators of local and systemic inflammation (KONNO et al, 2012), since they inhibit phagocytosis of zymosan particles and Pb yeast by macrophages. Finally, *in-silico* proteolysis studies with gp43 revealed that aggregation-prone, P10-like peptides could be generated by the action of several enzymes including proteinase K, trypsin and pepsin. Besides host endogenous proteases, the levels of proteases secreted by Pb including an aspartyl proteinase (pepsin-like) are increased upon infection (LACERDA PIGOSSO et al 2017), so we envision that P10 or a P10-like peptide may be formed under physiological conditions, giving rise to amyloid fibrils. Altogether these data call attention to the fact that candidate peptides for vaccine development can aggregate in solution with important implications for their efficacy and also safety.

## Materials and Methods

### Gp43 secondary-structure prediction

Gp43 secondary-structure prediction was performed using PSIPRED software (JONES, 1999, http://bioinf.cs.ucl.ac.uk/psipred/). PSIPRED is an accurate secondary-structure prediction method that uses BLAST (Basic Local Alignment Search Tool) to find regions of local similarity between known homologous sequences. Gp43 primary sequence (GenBank: AAG36697.1) was used as input for the analyses using standard parameters and the webserver retrieved as output the most probable conformation (α-helix, β-sheet or random coil) of each residue in the sequence (**Figure 1A)**.

### Prediction of solvent accessibility

The solvent accessibility of gp43 residues was predicted by NetSurfP server, available at: http://www.cbs.dtu.dk/services/NetSurfP-2.0/. NetSurfP is a web-based server with a user-friendly interface able to predict solvent accessibility based on sequence. It uses an architecture composed of convolutional and long short-term memory neural networks trained on large data sets of solved protein structures available on public data banks. The gp43 sequence in FASTA format was used as input to the prediction using default parameters.

### Calculation of GRAVY (Grand Average of Hydropathicity)

The GRAVY of peptides was calculated using the Expasy ProtParam toolkit available at: https://web.expasy.org/protparam/. ProtParam computes various physicochemical properties that can be deduced from a protein sequence. No additional information is required about the protein under consideration. Peptide sequences in FASTA format were used as input for the calculations. The GRAVY value for a peptide or protein is calculated as the sum of hydropathy values of all the amino acids, divided by the number of residues in the sequence.

### Generation of gp43 3D structural model

The gp43 3D structural model was generated by submitting gp43 primary sequence to RaptorX webserver (KÄLLBERG et al., 2012; http://raptorx.uchicago.edu/). RaptorX is a protein structure prediction server that excels at predicting 3D structures for protein sequences, when protein homologs with homology degree >30 % are available in the Protein Data Bank (PDB). RaptorX allows for template-based tertiary structure modeling and delivers high-quality structural models. As the template for the modeling, the high-resolution structure of β-glucanase (sequence similarity > 80 %) (PDB1CZ1) was used. The PDB file output from RaptorX was visualized, edited and colored with PyMOL (The PyMOL Molecular Graphics System, Version 2.0 Schrödinger, LLC) to generate the models presented in **Figure 1B**.

### *In-silico* analyses of aggregation propensity and the identification of amyloid-prone regions (APRs) in gp43

The intrinsic aggregation and amyloid formation propensities were evaluated either in the context of the complete gp43 protein sequence or considering the isolated derived peptides. These analyses were performed with AGGRESCAN (BELLI; RAMAZZOTTI; CHITI, 2011; SÁNCHEZ DE GROOT et al., 2005), employing default settings. ZipperDB (NELSON et al., 2005;THOMPSON et al., 2006) was used to detect the presence of APRs capable of forming steric zippers in the sequences of P4, P10 and P23 peptides, also employing default settings. Primary sequence of gp43 P10-like structure from *Paracoccidioides lutzii* glucan 1,3-β-glucosidase also used in the analysis was obtained from UNIPROT code C1H4T0.

### *In-silico* models of amyloid fibril cores

The recently developed program Cordax (LOUROS et al., 2020); https://cordax.switchlab.org/) was used to generate models of the amyloid cores (as steric zippers). The sequences of P4, P10 and P23 were analyzed using default settings. Cordax uses a large set of amyloid-core structures with atomic resolution deposited in the PDB as templates for the modeling approach. Briefly, to generate models of a query sequence, the program divides it into hexapeptides as the unit for the predictions (as in previously developed prediction methods). The side chains of the hexapeptides are modeled into all template structures of the library using the FoldX force field (GUEROIS; NIELSEN; SERRANO, 2002). FoldX yields a model and an associated free-energy estimate of the fitting (ΔG, kcal/mol). Cordax gives two main outputs: i. the prediction of whether or not the segment is an amyloid-core sequence and ii. the most likely amyloid-core model of that segment (the structural topology, orientation of β-strands and overall architecture of the resulting putative fibril core). The output of PDB files from Cordax were visualized, edited and colored to generate the structures in **Figure 3** using PyMOL.

### Gp43 *in-silico* digestion by proteases

*In-silico* gp43 digestion was performed using ExpasyPeptideCutter (GASTEIGER et al., 2005) to simulate the generation of cleavage products by different proteases. Enzymes predicted products of cleavage generated by the software were used to generate the data described in **Table 1** as P10-like peptides, namely, peptides which include P10 sequence added or deleted by few residues at the C-or N-terminals.

**Table 1.**
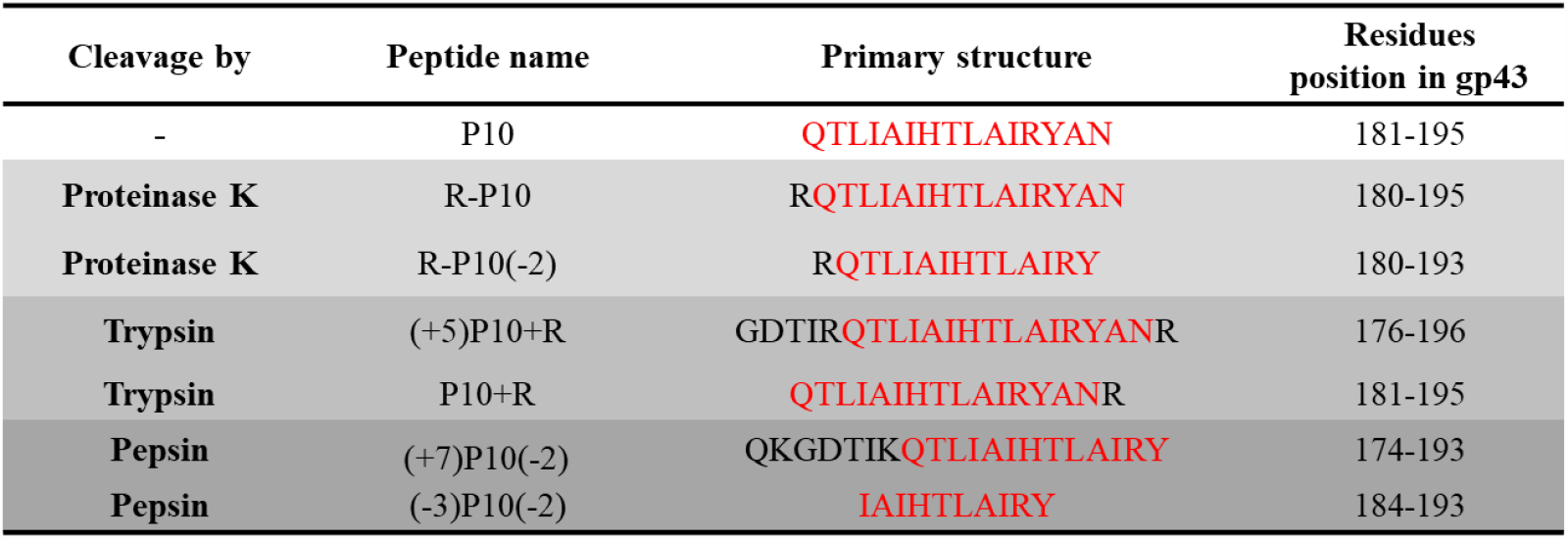
*In-silico* enzymatic digestion of gp43 shows that P10-like peptides can be formed. Full gp43 primary sequence was subjected to the ExPASy PeptideCutter tool to identify possible cleavage sites. Proteinase K, trypsin and pepsin were the enzymes predicted to cleave gp43, generating P10-like peptides while conserving P10 core sequence. The peptide sequences and their positions in gp43 are depicted in the Table, where: R-P10 = P10 with an extra R at the N-terminal; R-P10(−2) = the same as R-P10 shortened two residues at the C-terminal; (+5)P10-R = P10 with five extra residues at the N-terminal and R at the C-terminal; P10-R = P10 with an extra R at the C-terminal; (+7)P10(−2) = P10 with seven residues at the N-terminal and shortened two residues at the C-terminal, and (−3)P10(−2) = P10 shorted three residues at the N-terminal and shortened two residues at the C-terminal. In all sequences, P10 is colored in red.

### Peptides

Peptides purchased from BIOMATIK with purity above 95% were P4 (NLGRDAKRHLSKHWDTFITEDDFKNIAAAGL), P10 (QTLIAIHTLAIRYAN) and P23 (AFEVGAGWYFWTWKTEGAPGWDMQD). These were diluted in DMSO 100 % to a final concentration of 5 mM (stock solution).

### Circular dichroism (CD) assays

Peptide samples were prepared by diluting stock solutions immediately before measurements to a concentration of 100 μM in 2,2,2-trifluoroethanol (TFE) 5% (soluble samples) or in PBS (aggregated samples). CD spectra were recorded at the indicated times of incubation, scanning from 260 to 190 nm on a Jasco 810 spectropolarimeter thermostatted at 25 °C. Each spectrum shown is the accumulation of 10 scans.

### *In-vitro* aggregation assays

For *in-vitro* aggregation, P4, P10 and P23 stock solutions were diluted in PBS under agitation in the presence of Thioflavin-T (ThT; 50 μM), a specific fluorescent probe for amyloid fibrils. ThT fluorescence emission was measured at 485 nm by exciting the samples at 450 nm. To measure light scattering (LS), samples were excited at 320 nm while emission was collected at 320 nm. For the seeding assays, a solution containing 100 µM previously formed P10 amyloid fibrils was sonicated (40 KHz) during 30 min for mechanical fibril fragmentation. Five μL of a seed suspension were added to a freshly prepared P4 peptide solution at 100 μM under agitation. Seeds were used at the final concentration of 5 % unless otherwise stated.

### Mass spectrometry

P4-amyloid fibrils (formed in the presence of 5% P10 seeds) were centrifuged at 21,952x*g* for 30 min. After centrifugation, the supernatant was removed carefully, and the pellet was resuspended in 50 μL of a 9 M urea solution and incubated overnight under agitation. After that, the sample was diluted to 1 μM in 3% acetonitrile Wnano Acquity system (Waters, Milford, MA) for mass spectrometry. Proteins were desalted online using a trap column (Waters Symmetry C18, 180 μm x 20 mm, 5 μm) for 5 min and the liquid chromatography was performed with 3% to 85% acetonitrile containing 0.1% formic acid, 0.5 μL/min flow, in a HSS T3 130 C_18_ 100 μm x 100 mm, 1.7 μm analytical column (Waters, Milford, MA) for 58 min. System was set at initial conditions for 17 min in order to equilibrate the column.

Electrospray mass spectra were recorded using a Synapt HDMS quadrupole/orthogonal acceleration time-of-flight spectrometer (Waters, Milford, MA) interfaced to the nanoAcquity system. Capillary voltage was set at 3500 V, source temperature was 80 °C and cone voltage was 40 V. The instrument control and data acquisition were conducted by a MassLynx data system (Version 4.1, Waters), and experiments were performed by scanning mass-to-charge ratios (*m*/*z*) of 400–2000 using a scan time of 1 sec, applied during the whole chromatographic process. A 320 fmol GFP – Glu Fibrino Peptide solution in 50% acetonitrile containing 0.1% formic solution was set at a 0.5 μL/min flow and acquired during 1 sec after each 15 sec of the main chromatogram in order to calibrate spectra using Q-Tof’s LockSpray™ (Waters, Milford, MA).

All data were processed manually in MassLynx. To obtain an accurate molecular-mass measurement the resulting chromatographic peaks were analyzed, and the combined raw mass spectrum was lock-mass corrected in MS *m*/*z* scale using GFP ion 785.8426 m/z. Resulting spectra were treated using a charge-state deconvolution algorithm -Maximum entropy (MaxEnt 3, Waters, Milford, MA) -and monoisotopic singly charged ions were assigned in a relative intensity plot.

### Transmission electron microscopy (TEM)

Samples of aggregated peptides were diluted to 10 μM in Milli-Q water. Five μL of this suspension were absorbed onto 200-mesh carbon-coated copper grids for 5 min and then blotted to remove excess material. Negative staining was performed by adding 5 µL of 2% (w/v) uranyl acetate. Samples were dried in air for 3 min. The grids were imaged with a JEOL 1200 electron microscope (JEOL Ltd.) operating at a 60 kV acceleration voltage.

## Results

### Structural model and prediction of aggregation propensity of gp43 and its derived peptides

**Figure 1A** shows the primary sequence of gp43 (416 amino acids), highlighting in colored boxes the positions of the three peptides studied here, namely, P4 (green box), P10 (red box) and P23 (blue box). Since the tridimensional structure of gp43 has not been solved yet, its elements of secondary structure were predicted by using the program PSIPRED (JONES, 1999) and are depicted in this panel. The protein is predicted to be mainly composed of α-helices (13; marked in pink) with small β-strands (8; marked in yellow) and random-coiled/unstructured regions (marked in grey) intercalated among them. Gp43 has a primary sequence with high similarity to yeast β-glucanase (glucan 1,3-β-glucosidase). Since the high-resolution structure of β-glucanase has already been solved (PDB1CZ1) (CUTFIELD et al., 1999), we ran PSIPRED with β-glucanase as a control and the algorithm retrieved with great accuracy the known pattern of secondary structure elements of the enzyme (prediction: 14 α-helices and 8 β-strands; x-ray crystallography: 15 α-helices and 8 β-strands).

In the absence of a 3D structure of gp43, we built an *in-silico* model with RaptorX (KÄLLBERG et al., 2012; MA et al., 2012) using the yeast β-glucanase structure as template as previously done by other groups (KÄLLBERG et al., 2012; LEITÃO JUNIOR et al., 2014). **Figure 1B** displays the model of gp43 superposed on the structure of β-glucanase showing small deviations, most of them in loops/unstructured regions. **Figure 1C** shows in the generated model of gp43 the position of the three peptides studied here: P4 (green), P10 (red) and P23 (blue). As seen (**Figure 1A** and **1C**), the 31 amino acids of P4 are predicted to span an α-helix-rich region of the protein (amino acids 76-106), as well as the 15 amino acids of P10 (amino acids 181-195). Peptide P23, with its 25 amino acids (amino acids 372-396), however, encompasses a region supposedly devoid of a defined secondary structure with a small β-strand in its middle. As seen in **panel C**, while P4 and P10 are located on the surface, P23 is mainly buried in the core of the protein.

As mentioned before, these three peptides from gp43 are highly immunogenic, especially P10, which has been envisioned as an element in the development of a vaccine against Pb (MAGALHÃES et al., 2011; DE AMORIM et al., 2013). As it is evident in the structural model of gp43 (**Figure 1B and C**) and according to the predictor of solvent accessibility NetSurfP (KLAUSEN et al., 2019), a little more than half (8 out of 15) residues of P10 are considered highly exposed. These residues are Q181, I184, H187, T188, I191, R192, A194 and N195. This high degree of exposure is potentially incompatible with the high hydrophobicity of the peptide. The Grand Average of Hydropathicity (GRAVY) of P10 is 0.607, while that of Aβ peptide, which partially spans the cell membrane and is involved in Alzheimer’s Disease (SOTO; CASTAÑO, 1996), is 0.205 (ProtParam Expasy; GASTEIGER et al., 2005). This latter feature led us to consider whether P10, when in solution, might have a tendency to aggregate, and this property should be evaluated before using it for vaccine development.

In order to sort out the aggregation propensity of the different segments of gp43, including the P10 region, Aggrescan was employed (CONCHILLO-SOLÉ et al., 2007). Aggrescan is a well-validated algorithm that allows the identification and evaluation of aggregation-prone regions (APRs) within proteins and peptides. The Aggrescan data analysis for full gp43 is depicted in **Figure 2A**.

**Figure 2.**
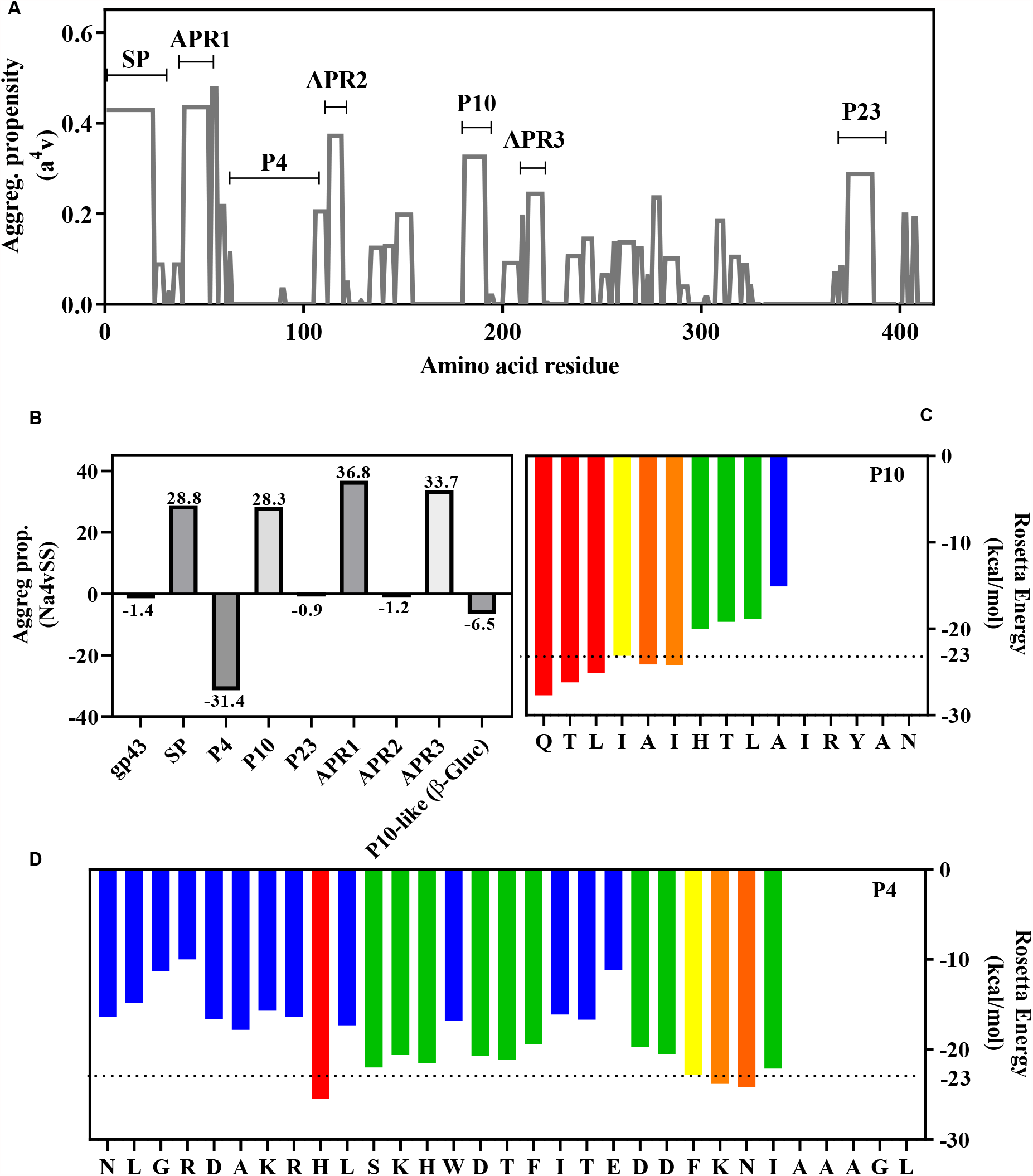
Aggregation propensity analyses of entire gp43 and its derived peptides. **(A)** Aggrescan analysis of entire gp43 showing its aggregation propensity scores along its primary sequence. SP: signal peptide (amino acids 1-35); APR1, 2 and 3: amyloid-prone regions 1, 2 and 3. Regions corresponding to P4, P10 and P23 are marked. **(B)** Aggrescan analyses of different segments of gp43 (SP, APR, P4, P10 and P23) as well as the P10-like segment in β-glucanase. (**C** and **D**) ZipperDB analyses of P10 and P4 showing hexapeptides predicted to form steric zippers (Rosetta design values bellow -23 kcal/mol). Yellow, orange and red bars indicate the first residue of a hit hexapeptide (N to C terminal); green and blue bars indicates that hexapeptides beginning on that residues do not fit into a steric zipper.

In **Figure 2A**, the regions corresponding to the three peptides of gp43 studied here are marked. As seen, P10 and P23 are inserted in regions with high aggregation propensity, while P4 is not. There are other regions dispersed along the protein with high aggregation propensities (those with high scores are labelled APR 1-3), including the region encompassing the signal peptide (SP) at the N-terminal region (∼ the first 50 residues).

**Figure 2B** depicts the aggregation propensities of the peptides under study, which are compared to the aggregation potential of the entire gp43 as well as with other APRs present in the protein. Overall, the whole gp43 protein has no tendency to aggregate. This is also true of P4 and P23, even though the latter spans a region of gp43 with high aggregation scores (**panel A**): when analyzed in isolation and normalized to its number of residues, it does not have a high aggregation score. On the other hand, P10 is among the APRs with higher aggregation propensity scores, like the signal peptide located at the N-terminal region of the protein (N-term 1-35). Interestingly, the region corresponding to P10 in β-glucanase (QTLAAI**R**ALAN**R**YA**K**) has no tendency to aggregate according to Aggrescan (**Figure 2B; P10-like**), probably due to the presence of positively charged residues (marked in bold), two in the middle (**R**7 and **R**12) and one at the end (**K**15) of the peptide, which might act as aggregational gatekeepers (MONSELLIER and CHITI, 2007; ROUSSEAU and SERRANO; SCHYMKOWITZ et al, 2006; SANT’ANNA et al., 2014). P10 in gp43 (QTLIAIHTLAI**R**YAN) has only one arginine (position 12). These observations have important implications and will be discussed below.

Next, ZipperDB (THOMPSON et al., 2006), an algorithm that predicts the ability of a peptide to form amyloid steric zippers, was employed with the three peptides of interest and the data are presented in **Figure 2C** (P10) **and D** (P4). The 3D-profiling method of ZipperDB was the first amyloid predictor built upon 3D structural data. At the core of amyloid fibrils is the cross-β spine, a long tape of β-sheets formed by the constituent protein or peptide (NELSON et al. 2005). In an amyloid steric zipper, the side chains from two β-strands form a tightly interdigitating dehydrated interface, so that the resulting β-sheet bilayer forms the fundamental building block of the fibrillar aggregates (NELSON et al. 2005). In the predictor approach, each hexapeptide from the query sequence is threaded onto the experimentally determined 3D-structure of the NNQQNY peptide, and the energetic fit is evaluated by using the RosettaDesign energy function (GOLDSCHMIDT et al., 2010).

According to the data presented in **Figure 2C**, in P10 there are several hexapeptides able to adopt an amyloid steric zipper structure, as evidenced by consecutive hot-colored bars (yellow-red) in its N-terminal region, where it reaches the threshold value of -23 kcal/mol. In the graph, the Rosetta energy value of each bar corresponds to the hexapeptide beginning at that position and running +5 residues to the right. P4 also presents 4 non-contiguous hexapeptides able to form zippers, three of them in its C-terminus and one in the middle of its sequence **(Figure 2D)**. P23 does not present any region in its structure able to be accommodated in a steric zipper structure (**not shown**).

Since ZipperDB pointed out the presence of several hexapeptides capable of forming amyloid steric zippers, we used the recently developed program Cordax (LOUROS et al., 2020) to build structural models of the putative amyloid zippers present in P10 (**Figure 3A**) and P4 (**Figure** 3B). Interestingly, Cordax detected the presence of 5 hexapeptides capable of self-interacting to form steric zippers as amyloid cores in the P10 sequence. As seen in the models presented in **Figure 3A** (**upper images**), TLAIRY, IAIHTL, LIAIHT, TLIAIH are supposed to form zippers composed of parallel β-sheets (sheets presented in yellow or red), while HTLAIR adopted an anti-parallel β-sheet topology. As mentioned before, in a steric zipper structure, the side chains of the residues in one strand placed in one side of the sheet face the side chains of residues in the strands belonging to the complementary sheet, creating the bilayer of the amyloid steric zipper. **Figure 3A** (**lower images**) presents the details of this side-chain complementarity observed in the zippers formed by P10.

**Figure 3.**
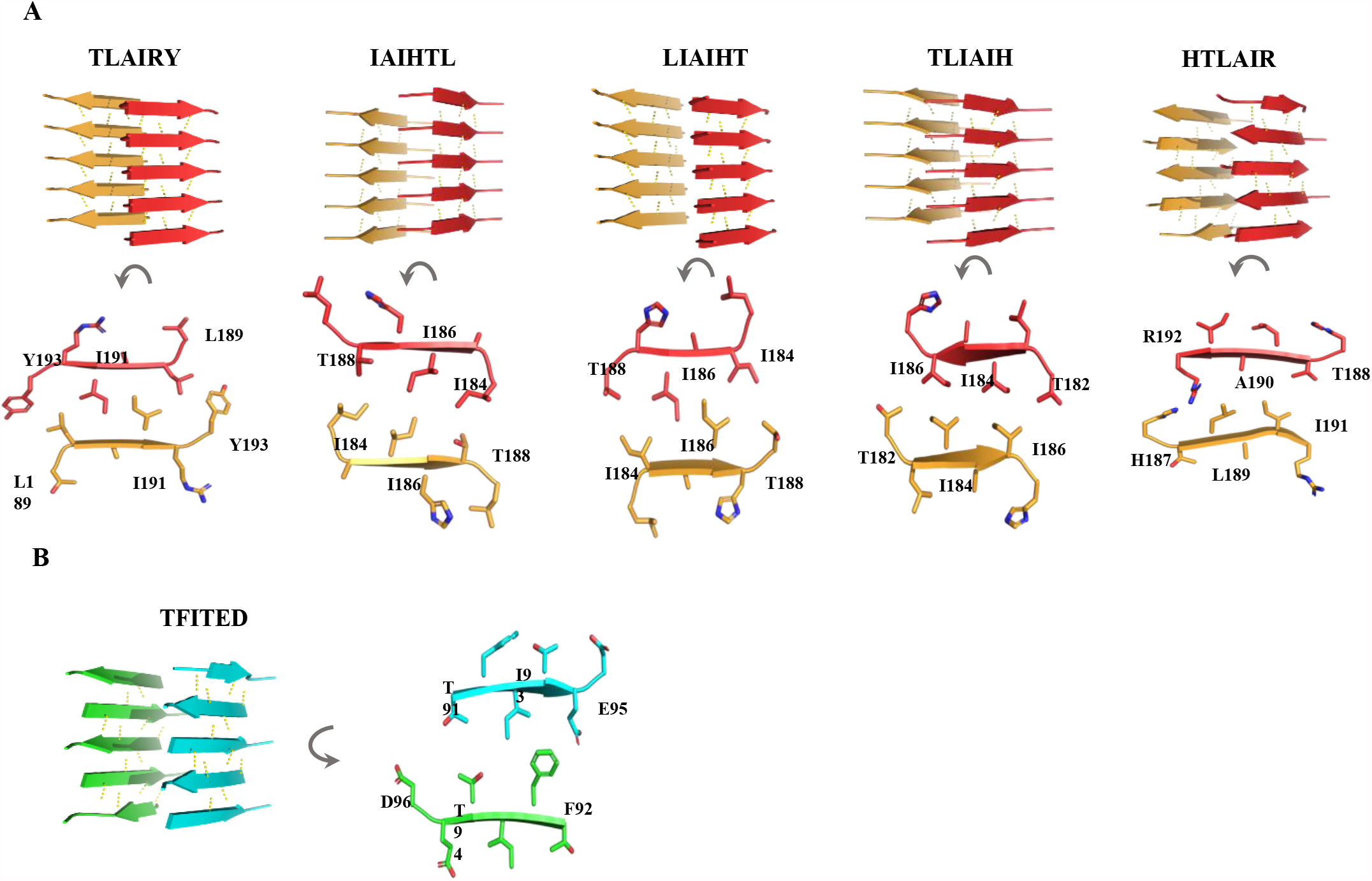
Hexapeptides derived from P10 form steric zippers *in silico*. P10 (**A**) and P4 (**B**) sequences were analyzed by Cordax (LOURO et al., 2020). The algorithm identified five hexapeptides in the sequence of P10 (TLIAIH, LIAIHT, IAIHTL, TLAIRY and HTLAIR) capable of forming steric zippers. Except for HTLAIR, each one forms two sheets (orange and red) composed of parallel β-strands (antiparallel in HTLAIR). In panel **A**, the lower images show details of the interdigitation of the lateral chains of the amino acids in the steric zippers facing the interior of the bilayer. (**B**) Only one hexapeptide of P4 (TFITED; blue and green) adopts a steric zipper structure in an antiparallel fashion.

Regarding P4, only a hexapeptide (TFITED) in the central region of the sequence is capable of forming an amyloid steric zipper (**Figure 3B**) in an anti-parallel architecture. As seen in the primary sequence of P4, out of 31 residues, 10 are charged (NLG**RD**A**KR**HLS**K**HW**D**<u>TFIT</u>**<u>ED</u>D**F**K**NIAAAGL, excluding histidine) and are dispersed along its sequence, and these charges probably hinder zipper formation. The two charges of TFIT**ED** are contiguous and present at the C-terminus of the peptide. As seen in the predicted model of this steric zipper (**panel B, right**), these two negative charges are in opposite direction in the zipper facing TF residues. In line with ZipperDB predictions, Cordax did not detect hexapeptides capable of forming amyloid steric zippers in the sequence of P23.

Taken together, these bioinformatics analyses indicate that P10, the peptide with the highest antigenic properties of gp43, presents high aggregation propensity, being able to form amyloid steric zippers at least *in silico*, a property that should be taken into account as it is a candidate for vaccine development. The next experiments were performed in order to unravel whether or not these peptides aggregate in solution.

### P10 forms amyloid fibrils in aqueous solution at neutral pH and seeds the aggregation of P4

Since *in silico* approaches indicate that at least P10 has APRs, *in-vitro* studies were carried out to evaluate their aggregation properties. In order to evaluate the secondary structures of P4, P10 and P23 in solution before aggregation, circular dichroism measurements were performed. The solvent TFE had to be used to keep the peptides soluble. As seen in **Figure 4A**, P10 and P4, when dissolved in TFE, assume α-helix-rich structures, while P23 has a spectrum more closely related to a β-sheet-rich peptide (**Figure 4A, inset**). This secondary structure data somehow recapitulates what has been predicted from the gp43 structure, when it was modeled onto the structure of β-glucanase (**Figure 1**). Accordingly, in this model, P10 and P4 encompass regions of the protein rich in α-helices, while P23 seems to be located in a region of the protein devoid of secondary-structure elements with only a small β-strand in it.

**Figure 4.**
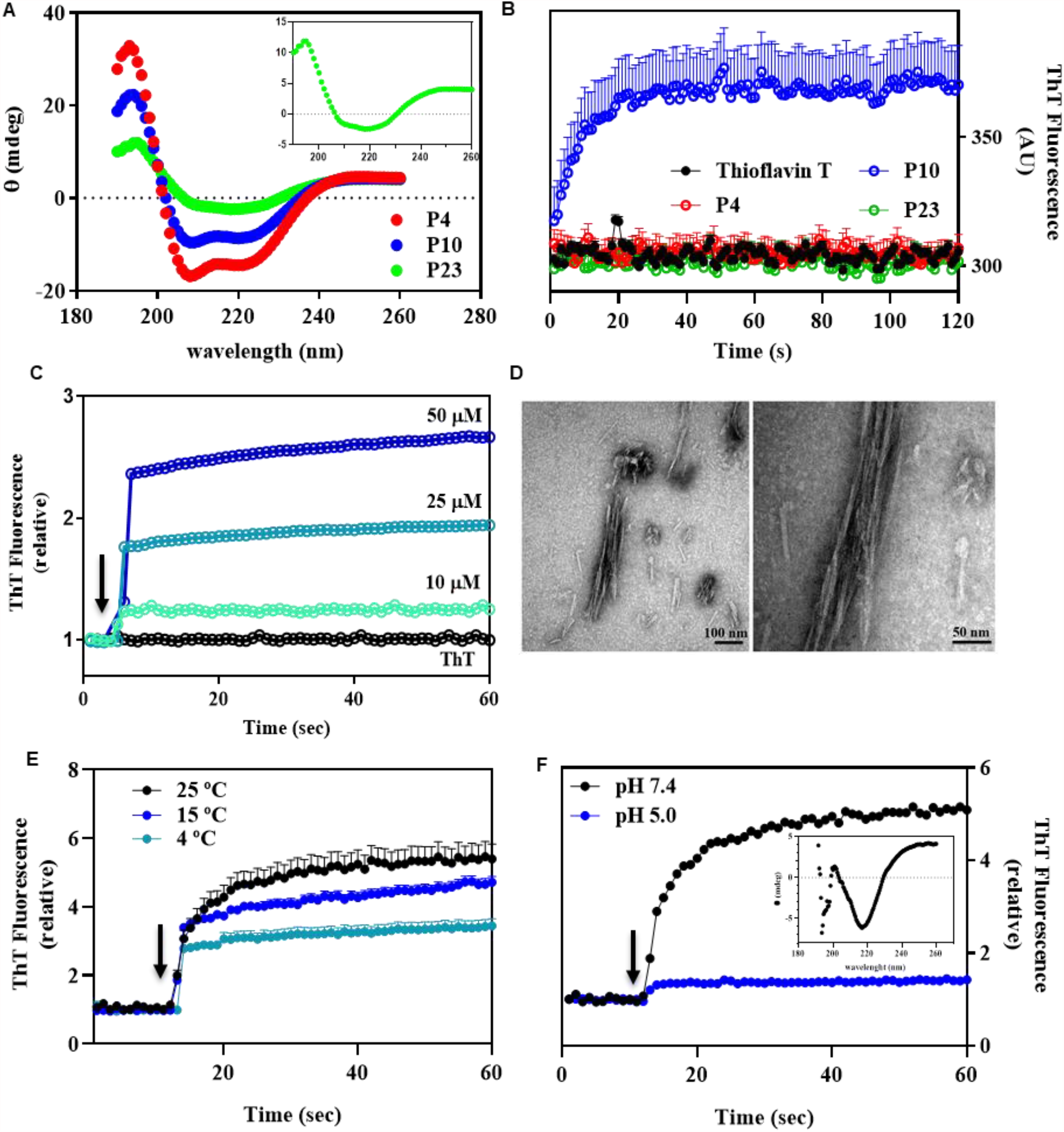
In solution, P10 undergoes aggregation forming amyloid fibrils. **(A)** Circular dichroism spectra of 100 μM of soluble P4, P10 and P23 in 5% TFE (see also the inset for more details of P23 secondary structure). **(B)** Kinetics of P10 aggregation (blue) at pH 7.4, 25 °C by measuring Thioflavin-T (ThT) binding. P4 and P23 do not aggregate in solution under these conditions. In all experiments (except 4C as indicated), the concentrations of peptides were 100 μM. (**C**) Aggregation of P10 (pH 7.4; 25 °C) shows concentration dependency. [P10] = 10, 25 and 50 μM, as indicated. (**D**) TEM images show the presence of mature amyloid fibrils composed of P10 formed at pH 7.4, 25 °C; right image is zoom of the left one. (**E**) Aggregation of P10 at pH 7.4 (100 μM) diminishes at low temperatures: Kinetics were performed at 4, 15 and 25°C, as indicated. (**F**) Aggregation of P10 at 25 °C, 100 μM at pH 5.0 (blue) and 7.4 (black). The arrows in panels C, E and F indicate the moment when the peptides were added to the buffer. ThT binding was measured by exciting the samples at 450 nm and recording emission at 485 nm. Error bars in (B) and (E) are SD in three independent experiments. (C), (E) and (F) error bars are shorter than the size of the symbol.

Next, the peptides (100 μM) were incubated in aqueous solution (pH 7.4, 25 ^°^C) in the presence of Thioflavin-T (Th-T), a specific probe for amyloid fibrils. **Figure 4B** shows that P10 forms Th-T-positive aggregates in ∼10 sec, which is suggestive of their amyloidal nature, while P4 and P23 did not. The next set of experiments was performed with P10, since this was the only peptide able to aggregate in solution, in agreement with most of the predictions (**Figures 2 and 3**). **Figure 4C** shows the aggregation profiles at pH 7.4, 25 ^°^C of P10 at 25, 50 and 100 μM as measured by Th-T and light scattering (**not shown**), confirming the amyloidal aggregation of this peptide in very short times, as well as its concentration dependency. Images of the aggregates formed are depicted in **panel D**, where it is possible to see mature amyloid fibrils, which present the typical β-sheet-rich structure measured by circular dichroism (**inset of panel F**).

**Figure 4E** shows that P10 amyloid aggregation is diminished at low temperatures (pH 7.4), as observed in other aggregation processes (SABATÉ et al., 2012).

Since the aggregation of proteins and peptides is influenced by pH and the endosomal compartments, where antigens are processed and docked at the MHC II cleft are acidic, we evaluated the aggregation of P10 at pH 5. **Figure 4F** shows that P10 behaves differently at pH 5.0 and its aggregation was almost completely abolished. These data are interesting and suggest that in endosomal compartments, aggregation might be prevented.

Next, we asked whether aggregation of P10 would occur in complete or incomplete Freund adjuvants, the vehicle used to produce vaccines. Interestingly, when dissolved in adjuvants at 100 μM, 25 ^°^C (pH ∼6.5), P10 did not undergo aggregation. Instead, it remained soluble, as measured by ThT binding (**Supplementary Figure 1**).

Amyloid formation can be accelerated by the presence of small fragments of mature amyloid fibrils, called seeds (HARPER; LANSBURY, 1997; JARRETT; LANSBURY, 1992). We asked whether seeds derived from P10 fibrils (sP10) would be able to seed the aggregation of P4 and P23. Interestingly, the addition of 5 % seeds of P10 to a solution with 100 μM P4 (pH 7.4, 25 ^°^C), which does not aggregate by itself, was able to induce immediately the aggregation of P4 as seen by Th-T binding (**Figure 5A, green curve**) and EM imaging (**Figure 5B; P4+sP10**). Similar experiments were tried with P23, but this peptide forms a viscous solution when diluted in aqueous buffer, which makes this type of experiment unfeasible. This result suggests that seeds composed of P10 catalyze the aggregation of P4, even though P4 does not have a tendency to aggregate in solution. It may be that the steric-zipper architecture predicted for P10 fibrils might catalyze the aggregation of P4 through the unique region of P4 with propensity to form zippers (TFITED; **Figure 3B**).

**Figure 5.**
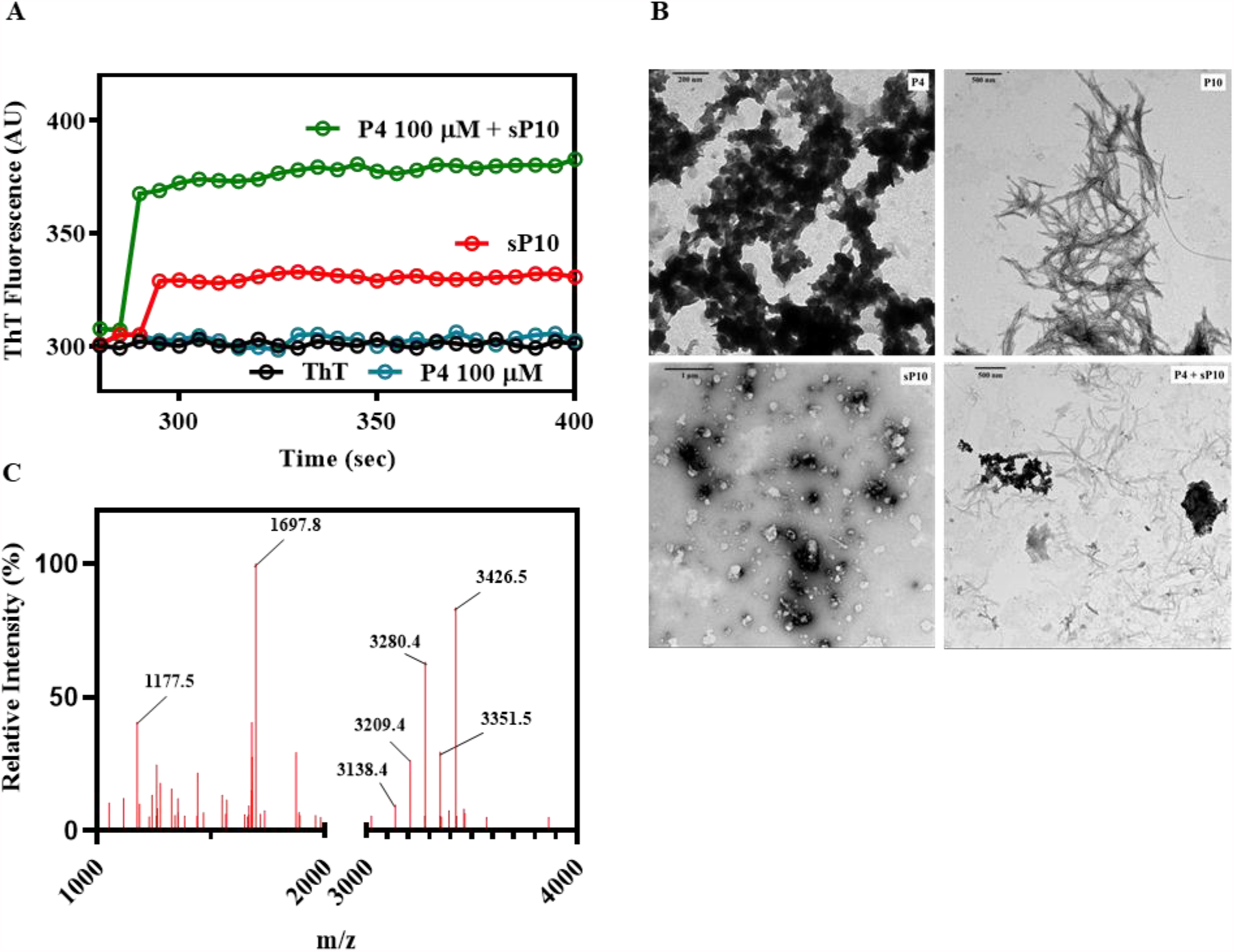
Seeds composed of P10 (sP10, 5%) are able to seed P4 amyloidal aggregation. **(A)** P4 alone (blue, 100 μM) does not aggregate in solution unless P10 seeds are also present (green). In red is shown the ThT signal of the seeds alone in solution and in black is the ThT emission when free in solution. **(B)** TEM images of the amorphous aggregates formed by P4 alone (P4, upper left), P10 fibrils (P10, upper right), seeds of P10 (sP10; lower left) and aggregates formed when P4 is seeded by P10 (P4+sP10; lower right). **(C)** Mass spectrometry analysis of the aggregates formed by P4 seeded with P10 showing the expected molecular masses of P10 (mass: 1697) and P4 (mass: 3411) suggesting that these fibrils grow by incorporating P4 into P10 seeds. Seeds of P10 were prepared as stated in Materials and Methods. All experiments were performed at pH 7.4, 25 °C.

In order to confirm that seeds of P10 were indeed seeding the aggregation of P4, amyloid fibrils were collected by centrifugation from the seeding experiments, washed, resuspended in 9 M urea for their complete dissociation and analyzed by mass spectrometry (**Figure 5C**). As seen, a peptide with the molecular mass of P4 (3,426.5) was present within these fibrils, as well as a peptide with the expected P10 mass (1,697.8). Other molecular masses detected were compatible with P4 and P10 degradation products (P4: 3,351.5; 3,280.4; 3,209.4 and 3,138.4; P10: 1,177.5)

### Aggregation propensity of P10-like peptides derived from gp43 proteolysis: *in-silico* analysis

Another important information to be taken into account is that gp43 is secreted from the yeast cells during infection, thereby being exposed to several proteases from the host. Besides, although in lower quantities then other secreted proteins, gp43, together with GAPDH and aspartyl proteinase, are increased and secreted during yeast biofilm formation (SARDI et al, 2015). This latter protease may also target gp43, fragmenting this protein into small pieces. Thus, we examined whether gp43 could be processed by different proteases by using the program ExPASy PeptideCutter (GASTEIGER et al., 2005). **Table 1** presents the result of these analyses showing only the primary sequence of the peptides in which P10 or P10-like peptides are formed. As seen, proteinase K digestion of gp43 is able to release the peptides R-P10 (P10 with an extra R at the N-terminal) and R-P10(−2) (the same as R-P10 shortened by two residues at the C-terminal), while trypsin generates (+5)P10+R (P10 with five extra residues at the N-terminal and R at the C-terminal) and P10+R (P10 with an extra R at the C-terminal). Pepsin, which is an aspartic protease of the same family as Pb protease, also gave interesting results and the fragments generated by its enzymatic activity are (+7)P10(−2) (P10 with seven residues at the N-terminal shortened two residues at the C-terminal) and (−3)P10(−2) (P10 shortened three residues at the N-terminal and shortened two residues at the C-terminal). In all cases, the cleavage products contain the APRs present in the P10 sequence.

In **Figure 6** are presented the amyloidogenic propensities of all peptides derived from gp43 digested with proteinase K, trypsin and pepsin calculated by Aggrescan. These are compared with P10 and also Aβ 1-42 aggregation propensities. Aggrescan shows that P10 and all P10-like peptides derived from *in-silico* enzymatic digestion of gp43 exhibit substantial aggregation propensity scores, even higher than Aβ 1-42. Exceptions to this behavior are (+5)P10+R and (+7)P10(−2), which display much lower aggregation propensities. among them This indicates that the additional residues inserted at the N-terminal of P10 diminish its aggregation propensity, probably because these short extensions include charged residues, R, K and D, which display gatekeeper properties preventing their aggregation: (+5)P10+R contains one D and one R at its N-terminal extension, plus an additional R at the C-terminus, while (+7)P10(−2) contains two K and one D at its N-terminal extension.

**Figure 6.**
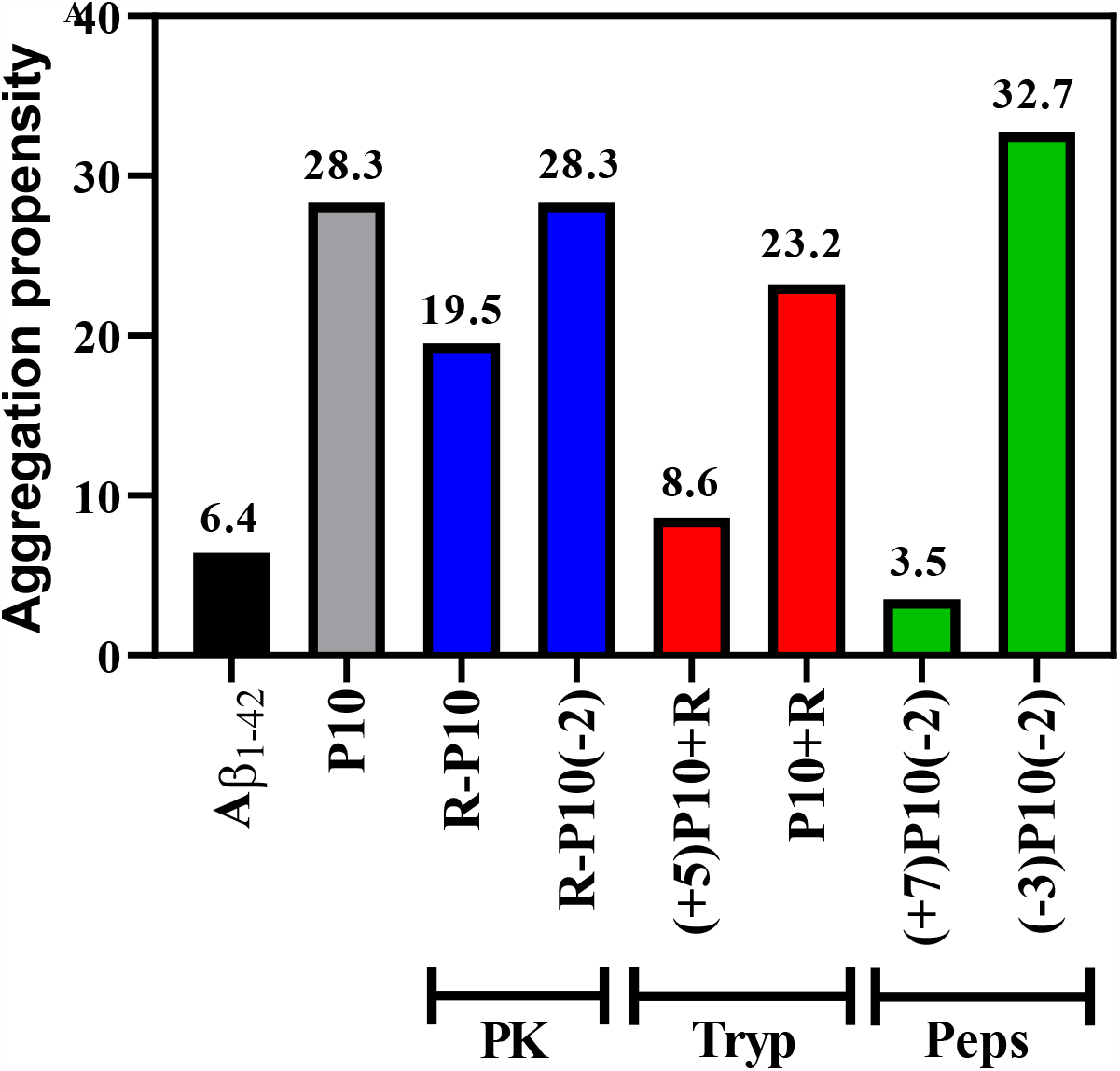
Aggrescan analyses of the peptides generated by *in-silico* enzymatic digestion of gp43 (see Table 1). **(A)** Aggregation propensity prediction by Aggrescan for each peptide generated by *in-silico* digestion of gp43 with proteinase K (PK), trypsin (Tryp) and pepsin (Peps). The aggregation propensity of Aβ 1-42 was included for comparison. For peptide nomenclature, see legend of Table 1.

Interestingly, the shortest of the peptides, namely (−3)P10(−2), generated by pepsin digestion, has the highest aggregation propensity among them all. This short peptide encompasses the sequence HTLAIR, the core of P10, carrying the essential region of the epitope (TRAVASSOS, 2004; MORAIS et al., 2000) which, as shown before (**Figure 3**), forms an anti-parallel steric zipper. Besides, it has only one charged residue in its extremity, which probably diminishes its gatekeeper property.

## Discussion

Peptide-based vaccines are promising, and several studies and clinical trials illustrate this application (reviewed by RAPPUOLI et al., 2016), including in COVID-19, where peptides derived from SARS-CoV-2 are being contemplated for this purpose (GRIFONI et al., 2020). Indeed, there are phase 1, 2 and 3 studies aiming to use peptides derived from different pathogens as vaccine candidates (LI et al., 2014).

Certainly, the initial search for a candidate peptide for vaccine development must be based on its immunogenic and physical-chemical properties. An initial possible strategy, among several others, is the use of one of several algorithms designed to help in the identification of putative epitopes in the primary sequence of a given protein capable of inducing a positive, desirable T-cell and B-cell immune response (RAPPUOLI et al., 2016). Additional evidence of the effectiveness of the identified epitope should be gathered, such as the identification of highly conserved immunodominant epitopes, the processing, presentation and association of the candidate peptide vaccine by antigen-presenting T cells in a highly MHC-heterogenous human population among others.

Once identified, these epitopes must be chemically synthesized as peptides for *in-vitro* and *in-vivo* studies, when its effective cellular and humoral responses must be evaluated as well as its protective potentials, when the host is challenged by the real pathogen. The choice of an adequate adjuvant is another important aspect of the process, due to the low capacity of peptides to induce innate immunity and a specific adaptative immune response, when administered alone.

However, the physicochemical properties of a candidate peptide must also be considered. Among them are its aggregation propensity, since aggregation could undermine a lot of the progress that has been made.

Here we used three peptides derived from the gp43 protein from Pb, one of which (P10) is considered as a candidate for vaccine development. They were chosen because previous studies indicated their immunogenic potential, especially P10. As shown here by different methodologies, P10 aggregates into amyloid fibrils when in solution at pH 7.4 (**Figure 4**). These fibrils exhibit the same architecture as that observed in amyloid fibrils present in amyloidogenic diseases (Eisenberg et al., 2012), but the other two peptides, P4 and P23, do not. However, aggregation of P4 was observed when seeded by P10 (**Figure 5**). Although adjuvants could change the aggregation behavior of P10 (**Supplementary Material 1**), we do not know what happens when the peptide in adjuvant reaches the plasma.

Bioinformatics studies with P10 indicate that there are regions within this peptide capable of forming steric zippers (**Figure 3**). According to a classification proposed by Eisenberg and co-workers (Eisenberg et al., 2012) in a detailed work where peptides derived from 14 diverse amyloidogenic proteins were shown to form zippers, these zippers can be classified into eight types of topologies. These topologies are distinguished by whether the strands in the sheets are parallel or antiparallel, whether the sheets pack with the same (‘face-to-face’) or different (‘face-to-back’) surfaces to one another forming the zipper, and whether the sheets are oriented parallel (‘up–up’) or antiparallel (‘up–down’) with respect to one another. Peptides TLIAIH, LIAIHT, IAIHTL, TLAIRY form type 1 zippers, where the strands are parallel in the same sheet, but the sheets are anti-parallel in respect to one another (up-down) and the sidechains of similar residues pair face-to-face (**Figure 3**). Interestingly, the zipper formed by the hexapeptide HTLAIR falls into class 7, where the strands on the sheets are antiparallel and they pack together face-to-back, where the zipper is formed by pairing of side-chains of different residues of the sequence (**Figure 3**). Regarding P4, it possesses only one hexapeptide stretch able to pack as a zipper, TFITED (**Figure 3B**). Interestingly, it adopts the same type of topology (class 7) as HTLAIR. These data are important because they model P10 into an amyloid fibril core, as shown by *in-vitro* experiments.

When in solution, P10 indeed forms mature amyloid fibrils, with the same architecture and secondary structure as those found in amyloid diseases as seen by TEM, ThT binding and circular dichroism (**Figure 4**). More intriguing is the fact that P10 fractured into seeds was able to seed the aggregation of P4 (**Figure 5**). There is some evidence that *Paracoccidioides brasiliensis* infection can trigger amyloid deposition in hamster kidney, affecting its physiological function, suggesting that this process can occur *in vivo* as well (FABRIS, 1976).

Another interesting property of P10 aggregation is its dependency on neutral pH (**Figure 4F**). There are interesting studies proposing that aggregation of antigens might be a strategy of phagocytic cells to concentrate and preserve the integrity of these antigenic peptides before their insertion into the MHC-II cleft and displacement of CLIP from the cleft (ZEPEDA-CERVANTES et al., 2018). Thus P10, when cleaved from gp43 during antigen processing in early/late endosomes (pH >5), might form aggregates inside these compartments until the pH is acidified by the fusion with lysosomes generating the endolysosomes (pH <5). This brings about the dissociation of the peptides from the aggregates in order to bind into the MHCII cleft, followed by migration to the cell membrane and presentation to another immune cell.

There are several examples of what has been called functional amyloids, such as the amyloid fibrils composed of hormones (ACTH, β-endorphin, GLP-2 and others), an efficient mechanism of storage and slow release. (MAJI et al., 2009). It may be that such a mechanism takes place during antigen processing and presentation, which might represent an efficacious immune strategy. Further studies with immune cells are necessary to confirm this hypothesis.

The tertiary structure of gp43 is not yet known, but due to its high sequence similarity with β-glucanase, it was possible to generate a homology model. P10 seems to be solvent-exposed in gp43, which makes sense since P10 is an effective epitope of Pb. Besides, a comparison between the primary sequence of P10 in gp43 (QTL**I**AI**HT**LA**I**RYA**N**) and that of the homologous region in β-glucanase (QTL**A**AI**<u>R</u>A**LA**N**RYA**<u>K</u>**) reveals an important feature (the amino acids that differ between them are in bold). Of special note is the presence of two additional charges (underlined, **<u>R</u>** and **<u>K</u>**), one in the middle and the other at the extreme of the C-terminal. Charged residues have gatekeepers’ properties that avoid the aggregation of peptides and proteins (ROUSSEAU et al. 2006; MONSELLIER and CHITI, 2007; SANT’ANNA et al., 2014). Interestingly, P10 from β-glucanase has no tendency to aggregate as evidenced by bioinformatic analyses (**Figure 2B**). However, P10 in β-glucanase lacks the HTLAIR sequence known to be the antigenic region (MORAIS et al., 2000; TRAVASSOS, 2004).

The sequence of P10 was established randomly by dividing gp43 into small peptides, which were further synthesized and tested for their immunogenic potential (TABORDA et al., 1998). P10 was extremely potent in inducing a consistent immune response in animal models (SILVA et al., 2017) by promoting a Th-1 lymphocyte response and the production of IFN-γ, also protecting the animals against further fungal infection (TABORDA et al., 1998). Later, using the algorithm TEPITOPE, which identifies immunodominant human T-cell epitopes of gp43 (NIELSEN et al., 2010), several different peptides were predicted to bind with high affinity to multiple HLA-DR molecules and P10 was the most promiscuous among them (IWAI et al, 2003), binding to 84% of the antigens. This suggests that gp43 could be processed by immune cells, which expose antigenic regions of gp43.

In order to get evidence regarding the formation of P10 or P10-like peptides by proteolysis of gp43, we took advantage of *in-silico* tools for prediction of protease cleavage products (**Table 1** and **Figure 6**). We agree that this is a first approach to gain insight into the formation of these peptides; however, we do not have gp43 purified to perform this investigation *in vitro*. Interestingly, even by using these *in-silico* analyses it was shown that proteinase K, trypsin and pepsin could digest gp43 forming “P10-like peptides”. Since gp43 is secreted by the fungi during growth, this protein encounter tissue-resident proteases, which can digest gp43 forming several peptides, among them the ones described in **Table 1**. Besides, Pb has an aspartyl protease similar to pepsin (TACCO et al., 2009), one of the proteases able to cleave gp43, generating P10-like peptides. These data suggest that these aggregation-prone fragments of gp43 may be formed under physiological conditions outside the cells. It is possible that fibrils of P10 and P10-like peptides contribute to Pb biofilm stabilization, as noted before in several other biofilms in which amyloid fibrils act as biofilm matrix scaffolds (SCHWARTZ et al., 2012; TAGLIALEGNA et al., 2016). Several of these fibrils are formed by proteolytic fragmentation of adhesins secreted by the pathogens (TAGLIALEGNA et al., 2016). Gp43 is an adhesin of Pb. Further studies are necessary to explore this possibility. We cannot forget that gp43 is processed by several resident proteases inside the endosome of immune cells before antigen presentation in MHC-II. During this process, P10 and P10-like peptides may be formed, and their high aggregation propensity has to be considered.

We also analyzed the aggregation propensities of the P10 like-peptides generated by the *in-silico* proteolysis. As seen in **Figure 6**, (+5)P10+R and (+7)P10(−2) have low aggregation propensities, as does P10 from β-glucanase, according to Aggrescan (**Figure 2B**).We can use this information to rationally design an antigenic P10-like peptide that retains the HTLAIR antigenic sequence, necessary for recognition of the MHCII cleft, but with low aggregation propensity, which could be useful in vaccine development. The proposed designed peptides are presented in **Table 2**. On the C-terminal end of P10 (QTLIAI<u>HTLAIR</u>**YAN**), the N should be replaced by K/R or a K/R should be added after N; on the N-terminal end (**QTLIAI**<u>HTLAIR</u>YAN) we propose the following alterations: QTLIAI should be preceded by additional charged residues, where the presence of three charges (two positives and one negative) seems to provide the best combination, as in the case of (+7)P10-2 (Q**K**G**D**TI**R**QTLIAI<u>HTLAIR</u>Y). It should be emphasized that the P10 position in β-glucanase has no charges to the left of its “HTLAIR” sequence, which appears in this enzyme as RALANR (QTLAAI**<u>R</u>**<u>ALAN</u>**<u>R</u>**YA**K)**. However, we note that this sequence in β-glucanase has an extra charge at its extreme N-terminal (R instead of H). This replacement probably decreases the aggregation potential of P10 in β-glucanase, but we cannot take advantage of this information to build our ideal peptide because it changes the antigenic region of P10 (HTLAIR). All these possibilities are presented in Table 2 with their aggregation propensity scores.

**Table 2:**
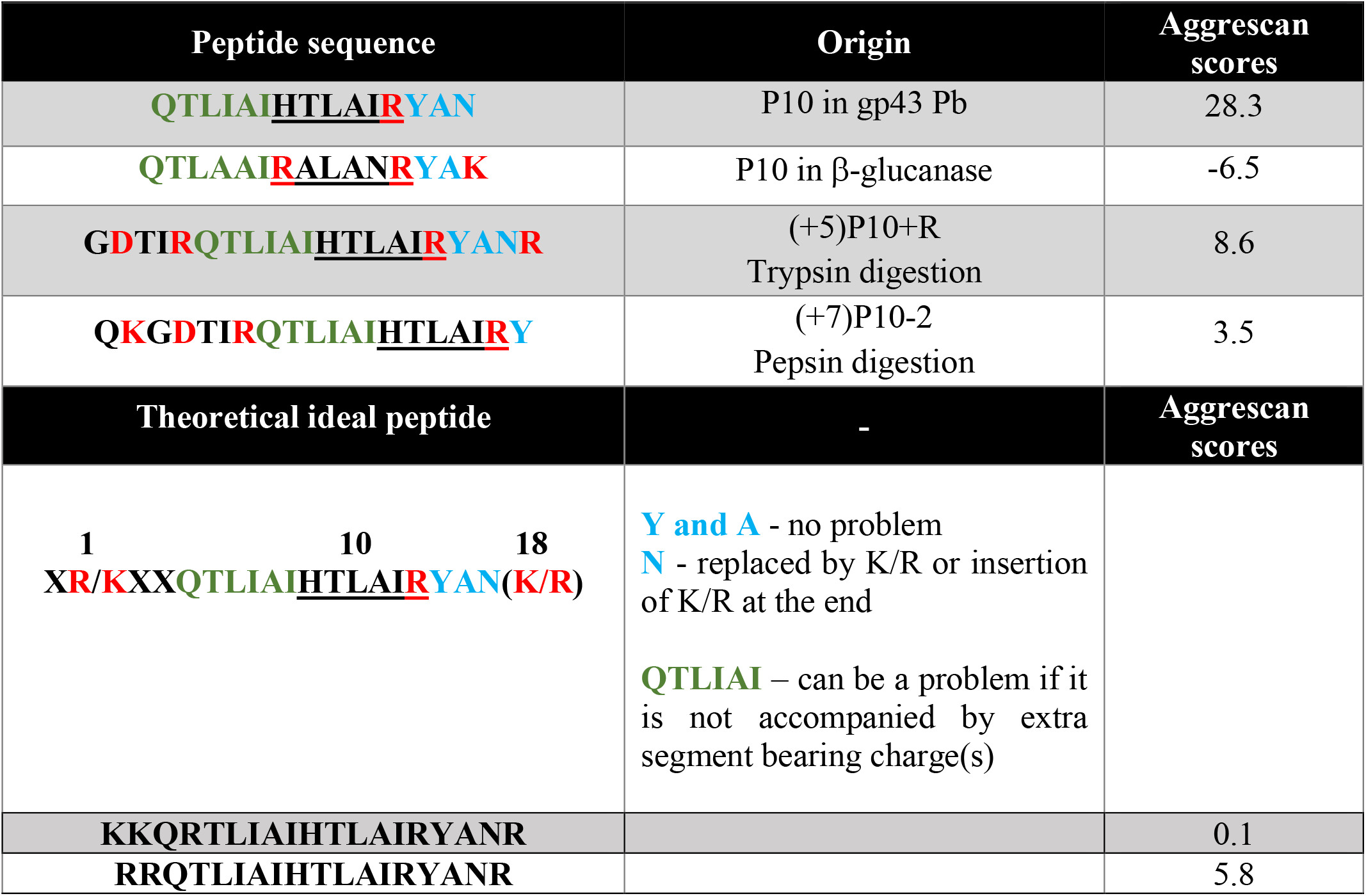
Rational design of a P10 ideal peptide for vaccine development. Charged residues are in red.

Taken together, this study aims to contribute to the design of candidate peptides for vaccine development, providing evidence for the need to study their aggregation propensity *in vitro* or even *in silico* to aid in the development of new effective vaccines. In addition, the possibility of obtaining aggregate-prone peptides derived from secreted pathogen proteins opens new possibilities for studying their functions in this aggregated state, a property of functional amyloids. More studies are necessary to unravel whether antigen processing in endosome compartments in the immune cells also takes advantage of the aggregation properties of antigenic peptides such as P10 for storage and release mechanisms.

## LEGENDS

**Supplementary 1 – P10 does not aggregate in vaccine adjuvants**. 100 μM of P10 were incubated in PBS (blue), complete (grey) or incomplete (red) Freund’s adjuvant containing thioflavin-T (ThT) to measure amyloid formation. **(B)** After aggregation kinetics, solutions were centrifuged and resuspended in a PBS-Thioflavin solution and ThT fluorescence emission spectra were recorded, confirming that fibrils were formed only in PBS. **(C)** P10 was aggregated at pH 7.5, 25 °C in the absence of ThT and after 60 min, ThT was added and its emission was recorded to evaluate fibril formation.

