## Supplementary figures and images for "Peptides derived from gp43, the most antigenic protein from *Paracoccidioides brasiliensis*, form amyloid fibrils *in vitro*: implications for vaccine development"

### Supplemental Material 1

#### Supplemental Material 1

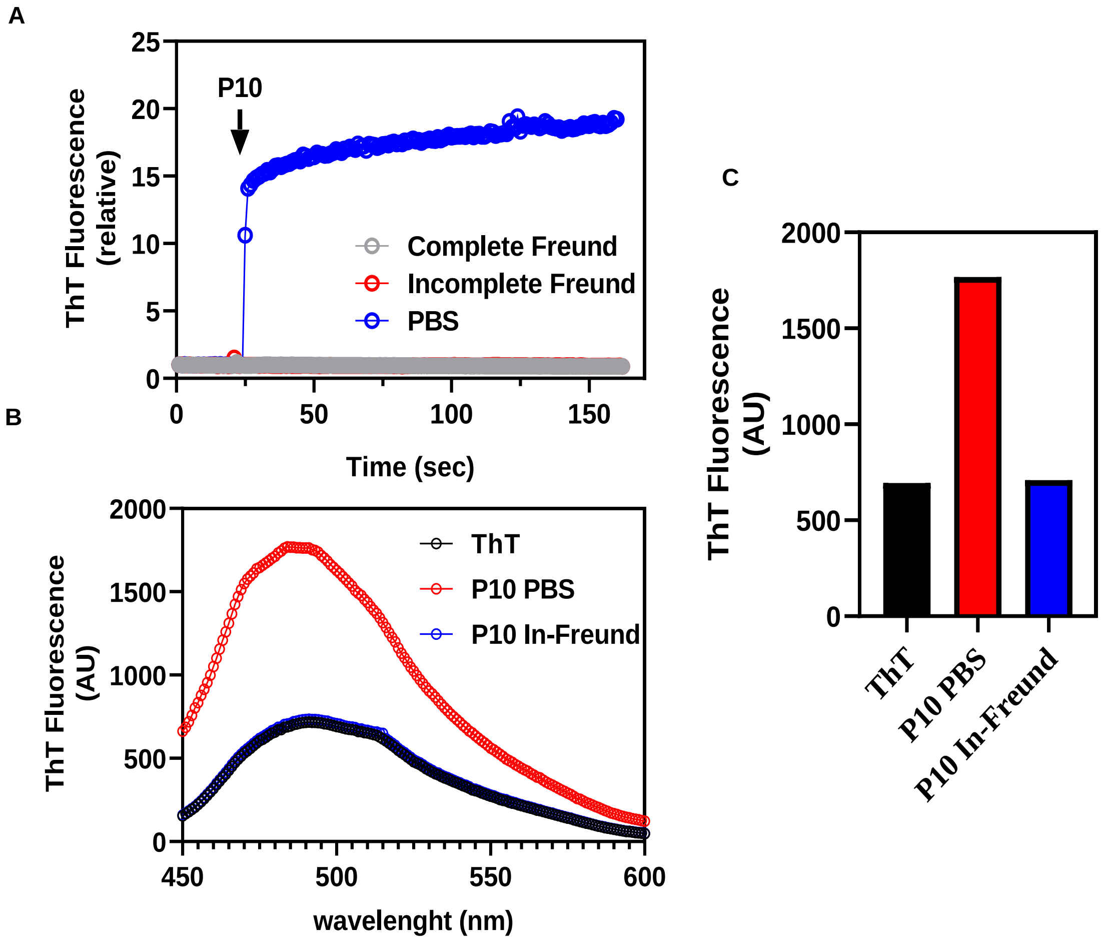
